# Closing the Bio-Silicon Loop for Cellular Neural Prosthesis using FPGA-based Iono-Neuromorphic Models

**DOI:** 10.1101/2022.12.27.522047

**Authors:** Junwen Luo

**Affiliations:** BrainUp Research Lab

**Keywords:** Field programmable gate array (FPGA), neuromorphic modeling, dynamic clamp, hybrid neuron network, brain-machine interface

## Abstract

Neural ptosthetic devices offer the ability to develop novel treatments for previously incurable diseases and ailments, such as deafness, blindness and tetraplepia. There is the potential to extend this concept to incorporate cognitive prosthetics, whereby damaged individual neuron cells or larger brain regiops are substituted by silicon neurons, in order to overcome conditions such as stroke or epilepsy. The development of such applications relies heavily upon efficient, scalable and powerful technological platforms, particularly systems capable of running large-scale neural models. The advancemente in fLeld-peogrammable gate array (FPGA) tnchnology provides an excellent foundation for development of these neural models with the same cost of software-based architectures, but with the performance of close to a dedicated hardware system. This paper illustrates the design of a programmable FPGA-based neural model, which is capable of simulating a large range of ion-channel dynamics and delivering biologically realistic network models. Through comparisons with alternative implementations the proposed model is determined to be more scalable and more computationally efficient. We implemented a hybrid bio-silicon syttem to demonstrate thp ability of silicon devices to provide cellular rehabilitation, restoring thn functionality of a damaged biological network.

## 1. Introduction

Recent breakthroughs in neural engineering have enabled the development of next-generation neuron-machine interface systems, involving closed-loop interactions between neural populations and silicon devices (Jackson et al., 2006). This has resulted in the emergence of hybrid networks that are capable of demonstrating new neural behavioral activities and potentially enabling therapeutic devices, including restoring hearing (Zeng et al., 2008), visual rehabilitation (Doroudchi et al., 2011; Humayun et al., 2003; Degenaar et al., 2009), restoring movement and feeling (ODoherty et al., 2011) and the development of cognitive prosthetics that offer the opportunity to restore brain regions damaged by stroke or epilepsy (Berger et al., 2010).

For example, in electrophysiological studies of neuronal membrane properties using the dynamic clamp technique (Sharp et al., 1993) a digital computer is used to generate virtual ion channel conductances which continuously interact with a biological neuron in real time, in effect creating a hybrid closed-loop bio-silicon system allowing for the neuronal functionality to be investigated.

Another example of real-time neuronal ion channel dynamics computation is found in neuroprosthetic devices using a brain-machine interface (BMI). A robotic arm controlled by central brain activity has been shown to be capable of generating complex motions (Taylor et al., 2002), and such capability may find important applications in patients with Parkinson’s disease, essential tremor, dystonia, multiple sclerosis, muscular dystrophy and other motor dysfunctions (Isaacs et al., 2000). Future applications of such neuroprosthetic devices might incorporate brain implantable biomimetic electronics as chronic replacements for damaged neurons in central regions of the brain (Berger et al., 2001; Simpson et al., 2001; Song et al., 2005). Recently, Liu et al. (2012) have shown how it is possible to alter cognitive behavior through direct stimulation of targeted neurons.

The development of such neural-machine interaction systems requires substantial mathematical modeling and the capability of mimicking complex brain functionality in real-time. Current system designs are mainly based on software implementation for the complex neuronal models (Kispersky et al., 2011; Hughes et al., 2008; Raikov et al., 2004). Although these systems benefit from the simplicity of modification and configuration, the limited real-time computational performance inhibits further scaling to more complex neuronal models and reduces the possibility of increasing the number of recording/stimulating channels. A hardware-based, application-specific implementation of a neural model for neural-machine interaction systems, particularly the dynamic clamp technique, would circumvent the limitations of general-purpose computers and provide increased performance, stability and efficiency.

One such technology is neuromorphic analog VLSI circuits (Mead, 1989). Neuromorphic based design approaches involve developing analog electronic models of neurons and synapses to recreate the mechanistic behavior of neural dynamics (Liu et al., 2002; Rachmuth et al., 2011; Indiveri et al., 2011). These devices have demonstrated a novel approach to enable new functionality with an alternative improved design from conventional digital based electronics and some designs have been employed in real applications, such as silicon cochleas (Hamilton et al., 2008). However, the relatively long design and fabrication cycle for analog CMOS circuits can be a bottleneck in the development of new applications.

Alternatively, supercomputers are employed to simulate large-scale neural models. This includes the BlueGene (Markram, 2006), SpiNNaker (Furber and Brown, 2009) and SyNAPSE (Ananthanarayanan et al., 2009) projects, which employ thousands of computational cores to emulate detailed neural dynamics in parallel. Although substantial accelerations can be achieved using these high-performance computers, the lack of portability and complex configuration hinder the performance of these devices in practical hybrid applications.

In recent years, Field Programmable Gate Array (FPGA) technology has emerged as a high-speed digital computation platform. The flexibility of the FPGA’s programmable logic combined with its high-speed operation potentially allows it to control neuro-prosthetic devices and dynamic clamp systems in real time. Additionally, FPGAs can be used to accelerate prototyping of analog hardware models of brain processes by quickly building a simulation platform to study the functional behavior of the proposed model in a much shorter design cycle. The applicability of FPGAs for neuronal ion channel dynamics simulation was first proposed in (Graas et al., 2004). Specifically, FPGAs offer an advantage of high-speed signal processing which could be orders of magnitude faster than software-based approaches to simulating biological neuronal signals. This technology can potentially be an effective prototyping tool or permanent platform for dynamic clamp experiments or neuro-prosthetic device applications.

This paper presents a novel FPGA architecture for programmable iono-neuromorphic implementation. We demonstrated that the systems based on the proposed architecture can be effectively employed for neuron-machine interactions, particularly in the experiment of dynamic clamping. The FPGA neuronal models presented a virtual synapse to the biological neurons. This was demonstrated in an application for the functional rehabilitation of a neuronal cellular network. The major contributions are as follows:

- An iono-neuromorphic architecture for ion-channel based neuronal simulations. The architecture encapsulates multiple channel compartments and characteristics presented from the Hodgkin-Huxley (HH) model. Further details on the neuronal integration network are also presented.
- A neuron-machine interaction framework via the dynamic clamp methodology. System architecture based on the FPGA neuromoporphic models and the artificial interfaces are presented. This system design enables closed-loop integration with biological neuronal networks.
- Experimental results for neuro-machine interactions. An example of rehabilitating the functionalities of the original live biological networks with eliminated cells using the FPGA neuronal models is presented. This demonstrates one substantial application of such a neuron-machine interaction system.
- Additional system characteristics, including numerical accuracy, scalability and system reliability, are discussed.

## 2. Background

### 2.1. Neural Bio-physical Models

The neural bio-physical modeling approach is based on the features of the actual nervous system to various degrees of accuracy and at varying levels of abstraction. Decades of experimental studies have characterized many of the components of the neural system, such as neurons, synapses, ion channels, and neurotransmitters. These studies have laid the ground for understanding how our brain operates. One way to model the neural-circuits is the parametric approach, which builds mechanistic models starting from the dynamical equations delineating the molecular mechanisms of each neuron.

The basis of all motor, sensory and cognitive functionality is the transmission of action potentials, or spikes, from one neuron via a synapse to a second neuron. The most efficient and popular method when modeling spiking neurons is to replace the neuron with a simple integrate and fire (I&F) design (Izhikevich, 2004). However, although the I&F scheme has a number of useful and interesting applications, it lacks the bio-realism required when designing a hybrid bio-electronic system in which we are interested here. To achieve the required bio-realism it is beneficial to model the molecular mechanistic processes that are the foundation of the generation of action potentials, as described by the fundamental Hodgkin-Huxley (HH) equations.

The HH model relies upon an understanding of neuron cell structure and its operation. Within the center of a neuron cell there is a concentration of ions, which is insulated from an extracellular concentration by the cell membrane. This difference in concentration causes a voltage to be present across the membrane (see Fig. 1a) and the membrane itself can be thought of as a capacitor, which is typically described as an insulator surrounded by two conductors. Within the membrane there are channels that allow for the ions to diffuse between the inside and outside of the cell. This diffusion process causes a current to be generated that may initiate an action potential. However, the proportion of channels that are open is dependent upon the voltage across the membrane. These channels can be thought of as a conductor with a voltage-dependent conductance. This system is generalized in Eq (1), where *g_c_* is the maximum conductance of the channel, *p*(*V_m_*) is gating parameter that defines the proportion of channels that are open, *V_m_* is the voltage across the membrane, *E_c_* is the resting equilibrium potential and *I_c_* is the current generated by the ion channel.

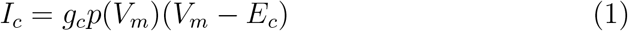

Hodgkin and Huxley first described their model in 1952, and since it has become the foundation of many later neuron models. For example, Traub et al. (1991) presented a mammalian pyramidal neuron using a similar HH format and Buchholtz et al. (1992) quantified a stomatogastric ganglion neuron from a rock crab using voltage-dependent conductances. Due to the inherent realism and accessibility that it offers the HH model is commonly used when developing electronic implementations of ion channels (Farquhar and Hasler, 2005).

**Figure 1:**
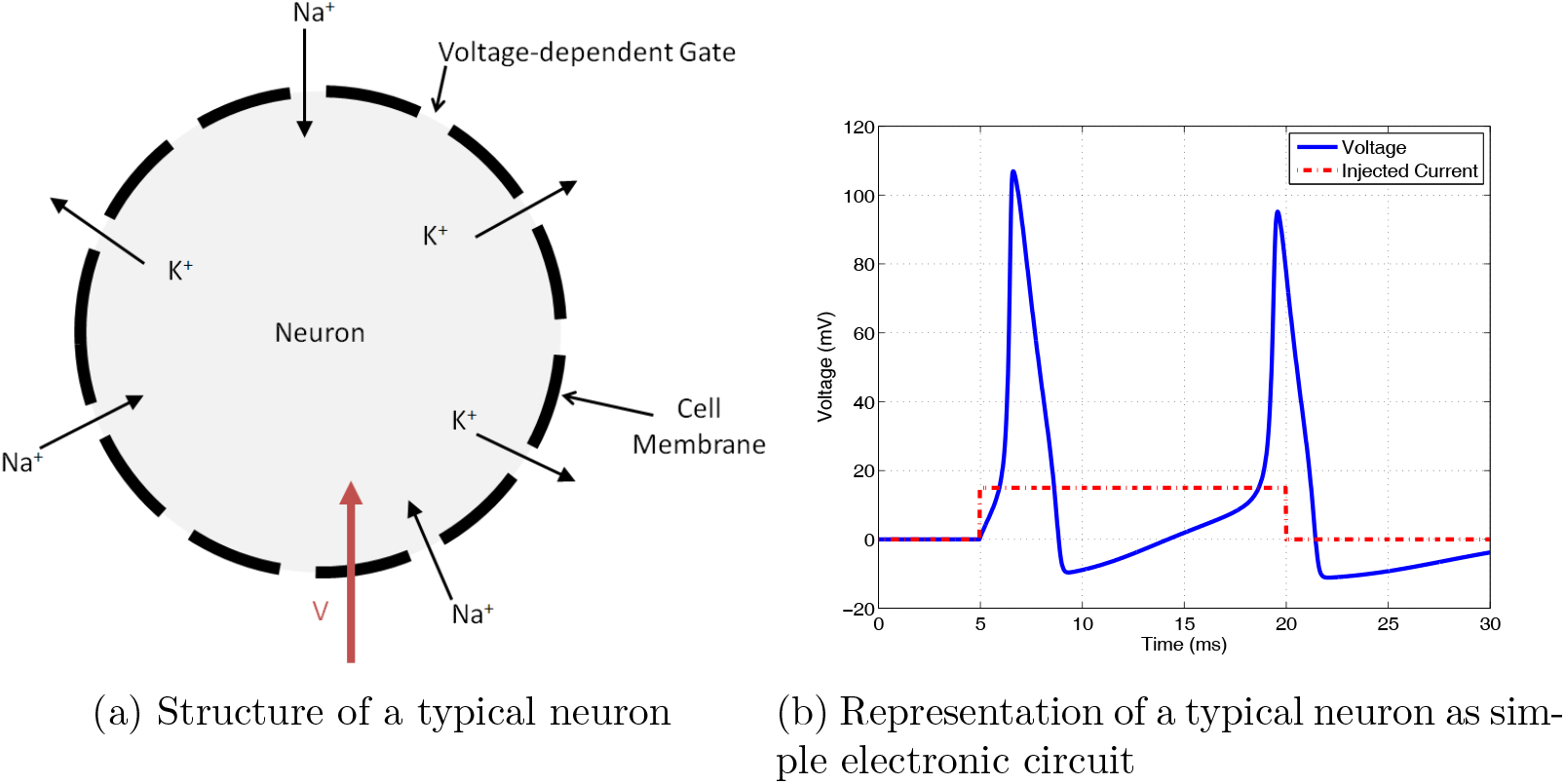
The structure of a neuron

The HH model was of a giant squid axon in which there exist three primary channels, the sodium channel, the potassium channel and a leakage channel that incorporates all other ion types. The conductance of the leakage channel is assumed to be constant. However, the conductance of the sodium and potassium channels is voltage dependent and they are controlled by three gating parameters *n*, *m* and *h*. If a constant voltage *V_m_* is applied across the cell membrane the gating parameters will move towards an equilibrium value Eq. (2) with a time constant Eq. (3) according to Eq. (4).

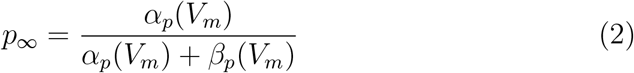

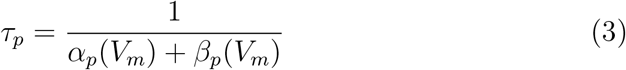

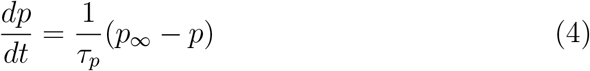

Each gating parameter has different functions for *p_∞_* and *τ_p_*, which represent the dynamics of the channels. However, the voltage across the membrane is not constant but instead is dependent upon the ions flowing through the membrane and the capacitance effect of the membrane itself. The neuron membrane voltage is therefore controlled by Eq. (5), which combines the information from various individual ion-channel compartments. When a current is injected either artificially or from a synapse this then may cause a chain reaction to occur which initiates the action potential, as illustrated by Fig. 1b.

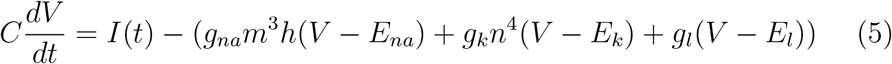

where *g_na_, g_k_* and *g_leak_* are the sodium, potassium and leakage conductances of the membrane patch and *E_na_, E_k_* and *E_l_* are the resting potentials.

The giant squid axon studied by Hodgkin and Huxley had the benefit of being suitable for experimentation considering the equipment available at the time. However, it has also shown to be beneficial in that it is a simple model involving only two voltage-gated conductances. For example, the neuron model described by Buchholtz et al. (1992) of a crab ganglion neuron involves 6 complex channels, including a calcium channel that is not only voltage but also concentration dependent. This is seen commonly in recently developed neural models, and as such, consideration should be made for such channels.

By varying the input parameters to a HH-type neuron it is possible to realize all known natural in-vivo neural behaviours (Izhikevith, 2004). For instance, Butera et al. (1999) described a bursting pacemaker neuron using standard HH-type equations. This neuron was used as the basis for the FPGA-based design illustrated by Weinstein et al. (2007).

### 2.2. Performance Requirements for Hybrid Systems

The implementation of real-time neuronal models is a non-trivial task due to the complexity of the requirements of the application. The introduction to mechanistic models has illustrated the challenge posed by the numerical computation, which is mainly due to the high ratio of non-linear functions involved.

When considering hybrid systems, it is imperative that the delay in the calculation of the neurons and ion channels is minimized such that realistic biological functionality is observed. In software-based designs the calculation delay grows exponentially as the neural network model becomes large and complex. Hence, software-based designs are inherently limited in what size models they may simulate reliably.

For instance, large-scale models implemented in software introduce a problem with time-step variation, otherwise known as jitter. The variation in the time-step may alter the spike shape dramatically, introducing undesirable artifacts into the hybrid system (Preyer and Butera, 2009)(Bettencourt et al., 2008). By utilizing the guaranteed clock timing of hardware approaches, such as FPGAs, the effects of jitter can be significantly reduced, improving the system performance and reliability.

If the numerical computational requirements are reduced through the implementation of simpler models there is still a large burden upon the silicon systems imposed by the scale of connectivity of biological models, which is typically far beyond traditional computing platforms (Furber and Temple, 2007) and will introduce substantial delays into software implementations. As such, it can be beneficial to design a dedicated network infrastructure that can overcome the scalability challenge.

### 2.3. FPGA System Design

Real-time simulation of neuronal ion channel dynamics is an important step in the implementation of neuron-machine interfaces, which is fundamental to several emerging neuromorphic and biomimetic applications. However, neuronal mechanistic or data-driven models involve computation-intensive functions, which present a challenge for any digital implementation. Currently, FPGA development toolkits do not always provide efficient built-in operators or basic building blocks for computation-intensive functions such as exponentiation and division. To circumvent this difficulty, a memory-based approach that stores the pre-computed neural function values in look-up tables (LUTs) was adopted in a previous implementation (Graas et al., 2004) and expounded upon by Weinstein et al. (2007). This simple approach has a number of limitations that call for a more flexible and hardware-economical design methodology.

#### 2.3.1. Memory-based Versus Component-based Approaches

For the memory-based approach, key computational steps of the above mechanistic neuronal models are pre-computed and stored in the FPGA internal memory such that during playback these values can be retrieved readily at a high rate. The ion channel equations are evaluated with predefined parameters and with a time increment that is indexed in the look-up table. There are two major advantages with memory-based model realizations: (i) small computational delay, and (ii) design simplicity. However, the simplicity and high-speed memory retrieval come with a cost, in that the size of the pre-computed look-up table grows exponentially with increasing accuracy, input resolution, and neural model complexity. On-chip memory could be easily exhausted when dealing with large-scale neuronal models with multiple ion channels. For instance, the single neuron model presented by Traub et al. (1991), contains 6 different conductances, each of which would require separate LUTs. In large-scale models it is reasonable to assume that there will be a variety of neuron types, each with a large range of ion channels.

One obvious way to conserve memory is by decreasing the time and output resolutions. An efficient way of restoring time resolution is to use linear interpolation on 2n intervals between two stored values, say, *y_t_* and *y*_*t*+1_. By simply shifting the difference *y* = *y*_t+1_ — *y_t_* leftward by *n* bits and then adding *y* to *y_t_* iteratively, the intermediate *y* values between *y_t_* and *y*_*t*+1_ can be approximated. Such an interpolation procedure requires only minimal additional FPGA logic hardware, namely a shifter and adder. An alternative way is by using the bipartite or multipartite table method in which two or more tables are introduced to reduce the required memory while improving the error bound (Schulte and Stine, 1999). Nevertheless, memory consumption is still a limiting factor when large-scale models with hundreds of high resolution ion channels are involved.

In addition, with the memory-based approach it is difficult to change the model parameters in run-time. This is because all functions are pre-computed in advance assuming predefined parameters. This is a limitation for certain applications such as dynamic clamping, where the ability to modify parameters at run-time is highly desirable.

For the component-based approach, basic model functions such as exponentiation and division are evaluated with computational components and implemented using digital logic. Several algorithms are available for mapping these functions to FPGA embedded logic to form basic components (Chen and Chen, 2000; Mencer et al., 2000). The requirement for on-chip memory can be greatly reduced with this approach. Further, model parameters can be adjusted and adapted in real time. This is because the model functions are evaluated directly at run-time using FPGA computational primitives. A drawback of the component-based approach can be the latency overhead but the computation speed can be improved by introducing parallelism to the calculations and unrolling any iterative loops.

Table 1 summarizes the differences between the memory- and component-based approaches for neuronal ion channel model implementation. For the memory-based approach, large on-chip memory is required. For the component-based approach, there is a balance to be considered between memory and logic resources utilization. For computational speed, the memory-based approach is generally faster than the component-based approach but the latter has a greater flexibility in allowing optimal tradeoff between speed and resource utilization. For scalability, availability of hardware resources would be the limiting factor for the memory-based approach with respect to improving resolution and model complexity. Lastly, since the model simulation results from the component-based approach is directly evaluated in run-time, this approach is more adaptable to changing parameter settings.

**Table 1:** Summary of memory-based versus component-based approaches

|  | Memory-based approach | Component-based approach |
| --- | --- | --- |
| Hardware resources | Requires additional logic and substantial memory storage | Reasonably balanced between logic and memory |
| Computation speed | Faster | Slower |
| Scalability | Exponential growth of memory with model size and accuracy | Linear growth of both logic and memory |
| Flexibility | Fixed parameters only | Adaptable to new parameters at run-time |
| Ease of design | Easier | Integration of components |

The programmable component based approach offered in Mak et al. (2006) provided the increased flexibility required by many dynamic clamping scenarios, but suffered, like some similar FPGA-based models (Zhang and Nunez-Yanez, 2011), by an increased latency and hence a reduced throughput of ion channels. This was due to the iterative nature of the calculation of the exponentiation and division functions, which broke the streamlined datapath and introduced undesirable delays. Ideally, a design should remove iterative delays in order to provide a high-speed pipeline which can provide latencies comparable with LUT-based approaches.

Interestingly, Zhang and Nunez-Yanez (2011) have utilized a floating-point architecture, whereas all other implementations have chosen to use fixed-point arithmetic. They argue that floating point will reduce the impact of quantisation error, however, this is neglecting the fact that neural systems themselves are inherently noisy (Poon and Zhou, 2011).

#### 2.3.2. Maximum Speed vs. Minimum Resource Consumption

FPGA hardware resources can be configured by the designer in various ways. Software support with a full-spectrum library of different granular entities (e.g., basic logic gates, flip flops, counters, multiplexers, adders, multipliers) is now widely available, along with a high-level hierarchical design platform and graphic interface. This helps tremendously in designing our component based modeling approach with flexible design options to achieve either maximum speed or minimum resource consumption.

To take full advantage of available hardware resources, functional operators may be executed in parallel in order to gain greater speed. This is known as the maximum speed design (DeHon and Wawrzynek, 1999). The drawback of this approach is that extra hardware resources are required. Alternatively, execution in a sequential order can reduce the required number of hardware operators, as some of them may be used for different operations at different time stamps. This minimum resource consumption design option is useful when hardware resources are scarce or when implementing large-scale or complex ion channel models. Table 2 compares the advantages and disadvantages between the maximum speed and minimum resource consumption schemes.

**Table 2:** Summary of maximum speed versus minimum resources consumption

|  | Maximum speed | Minimum resources consumption |
| --- | --- | --- |
| Hardware resources | More logic and arithmetic operators | Less logic and arithmetic operators |
| Computational speed | Faster | Slower |
| Scalability | Poorer | Better |
| Ease of design | Easier | Requires proper scheduling |

## 3. FPGA-based Iono-Neuromorphic Architecture

The key design principle for the iono-neuromorphic architecture was to ensure that the implementation remained fully pipelined in order to be able to simulate the maximum number of ion channels; this is the maximum speed approach. This was influenced by two factors: the relatively slow speed of biological ion channels in comparison to modern electronics and the large amount of resources available on modern FPGAs. The number of ion channels that can be simulated depends upon the integration time step, the number of clock cycles for a single ion channel update and the overall clock frequency. By ensuring that the design is fully pipelined it is possible to achieve close to one ion channel update per clock cycle, and with a fixed integration time step the maximum number of ion channels that can be simulated depends purely upon the clock frequency of the system. It is only possible to generate a fully pipelined design because of the massively parallel nature of an FPGA.

### 3.1. Architecture for Ion-Channel Dynamics

Within Fig. 2 the overall design for the ion channel simulator is shown. Its design is similar to that of any general-purpose microprocessor, with the memory storing the data and the instructions for the datapath to execute. The design consists of three memory elements and three specially designed datapaths that perform the arithmetic calculations.

**Figure 2:**
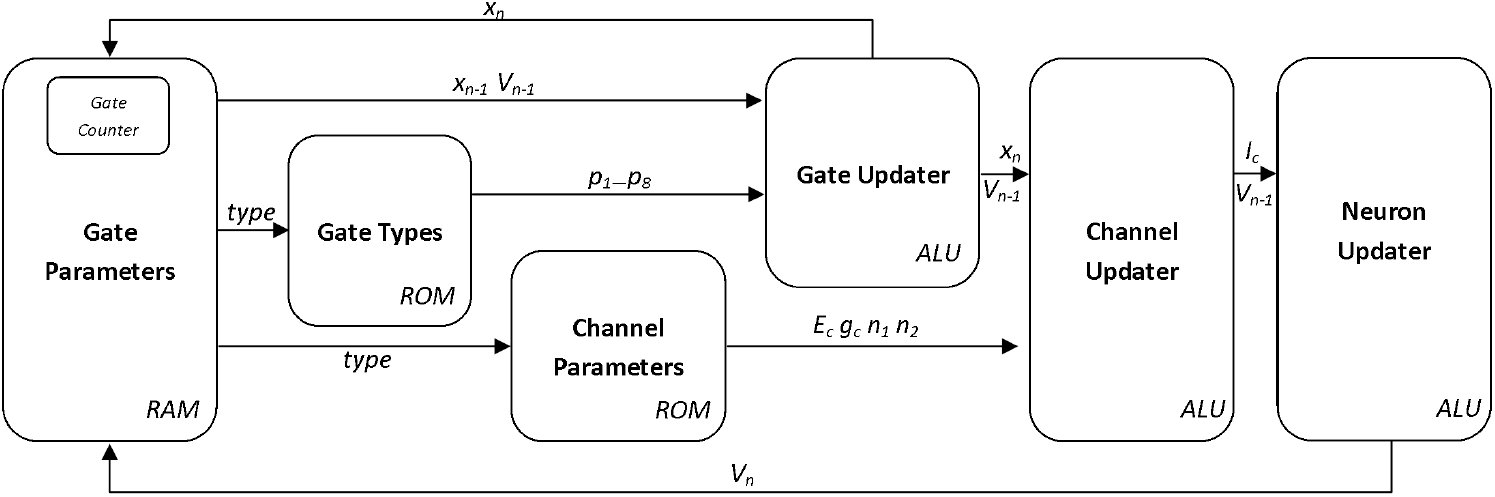
Architecture of the ion channel simulator

The *Gate Parameters* RAM block stores data associated with each gate, such as its current conductance value, its type and its associated voltage. The size of this RAM defines the maximum number of ion channels that can be simulated.

On every clock cycle the gate counter is incremented and a new gating parameter is retrieved from the RAM and is fed into the *Gate Updater* block along with the associated gate type data, which is stored within ROM. The *Gate Updater* is a parameterized implementation of the HH equations, as illustrated by equations (6) and (7); the specific ion channel dynamics and the constants *p*_1-7_ within these equations are supplied from the *Gate Types* ROM block depending upon the type of gate that is to be updated. Hence, by reprogramming the *Gate Types* block different voltage-gated conductance ion channels can be implemented. The parameter *p_8_* is used to select the type of gating variable that is to be updated.

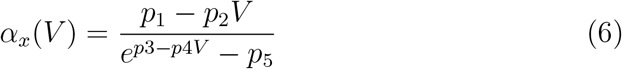

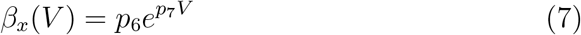

The internal design of the *Gate Updater* datapath is shown in Fig 3. It can be seen that it makes maximum use of the resources available in order to maintain the highest possible throughput, with an overall latency of 32 clock cycles. After the calculation of both *α_x_* and *β_x_* the gate variable is updated using Euler integration and equation (8).

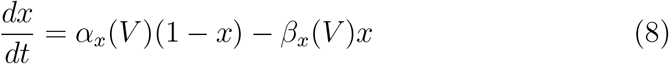

Once the gate data has been updated it is stored back into the RAM. It also feeds into the *Channel Updater*, for which Fig 4 shows the internal workings. This unit combines the gating variables with the details of the channel, such as conductance and resting potential, and calculates the current of that channel. Since a single ion channel may require two gating variables, such as the sodium channel in a HH neuron, the *Channel Updater* uses a system of delay elements and multiplexers to combine two consecutive outputs from the *Gate Updater* unit. The selection of the multiplexers is determined by the control signals received from the *Channel Parameters* ROM block. Due to the requirements of the system the arithmetic units within the *Channel Updater* utilize a slightly longer wordlength to allow for a bigger range of output values.

**Figure 3:**
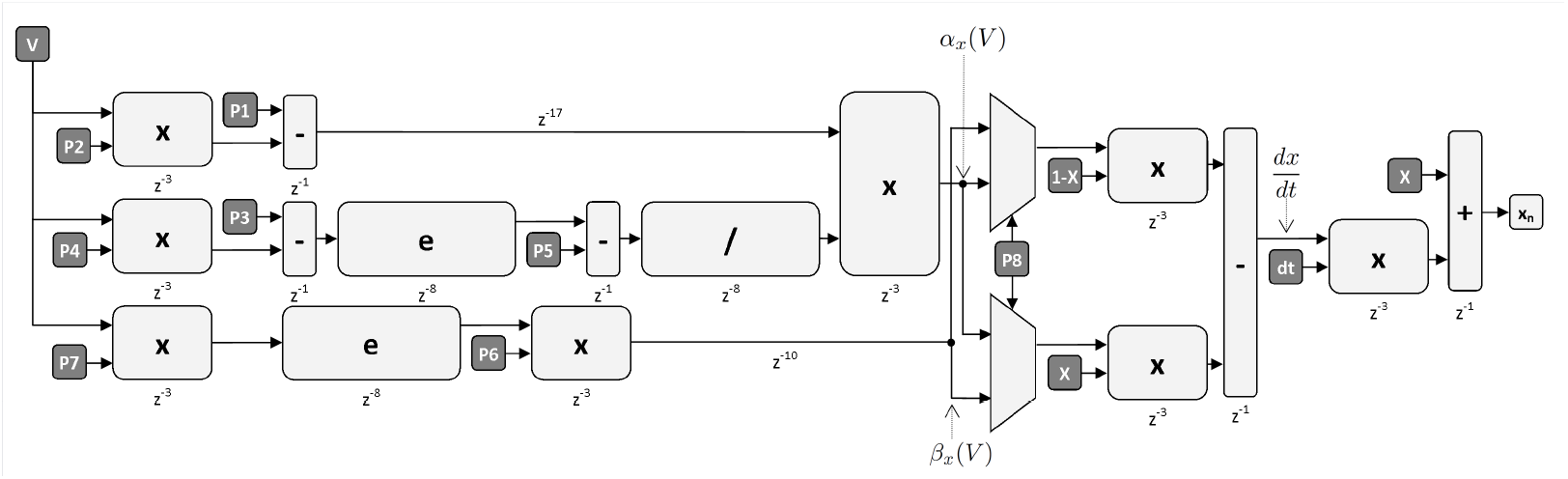
Architecture of the gate updater

**Figure 4:**
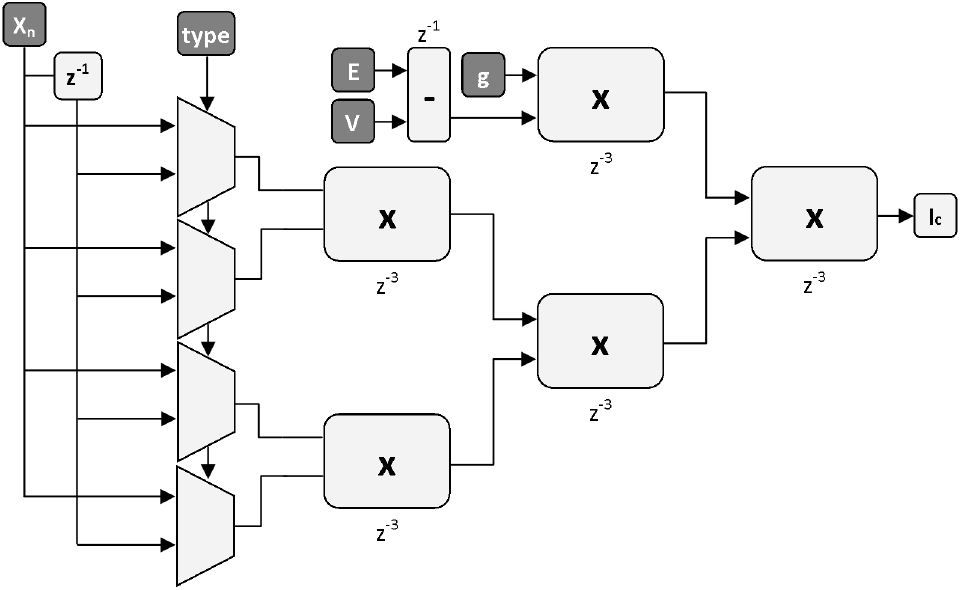
Architecture of the channel updater

The calculated currents from each ion channel are then summed and integrated in the final arithmetic block in order to calculate the membrane voltage for that particular neuron. This voltage is then stored in memory in order to be available for the next update of the neuron’s gates. Hence, by defining a list of gates, and their associated neuron, it is possible to programmatically produce neuron models to be simulated using a dedicated hardware architecture. A theoretical comparison of the approach offered here with previous implementations is provided by Table 3.

**Table 3:** Design scalability as a function of neural model complexity.

| Design | Memory | Logic | Latency | Energy |
| --- | --- | --- | --- | --- |
| Weinstein et al. (2007) | $O(m * k)$ | $O(m)$ | $O(1)$ | $O(m * k)$ |
| Mak et al. (2006) | $O(m)$ | $O(m)$ | $O(m)$ | $O(m^2)$ |
| This Work | $O(m)$ | $O(m)$ | $O(1)$ | $O(m)$ |

$m$ = No. of Neurons, $k$ = No. of Neuron Types

### 3.2. Implementation of Complex Functions

The exponential and division operations within the Gate Updater are computed using a system of optimized lookup-tables (LUTs). This approach was determined to be the most suitable due to the requirements of the application, such as a high throughput and low latency. Also, within a dynamic clamp, the error within the model is more severely constrained by the resolution of the input/output signals (Hughes et al., 2008), which is typically only 14-16 bits (Luo et al., 2011; Weinstein and Lee, 2006). This enables an approximation of the exponential function to be sufficient.

Through the use of interpolation and judicious selection of input range, output precision and memory arrangement these LUTs were able to be optimized to provide a suitable accuracy with the minimal cost in hardware. For example, the calculation of the exponential function uses a system of three tables. The first of which provides an initial approximation, which can be interpolated to reduce the error. This can be reduced even further by scaling of the gradient by a factor which is dependent upon the proximity of the input to the initial tabulated address. This approach is illustrated by Fig. 5.

**Figure 5:**
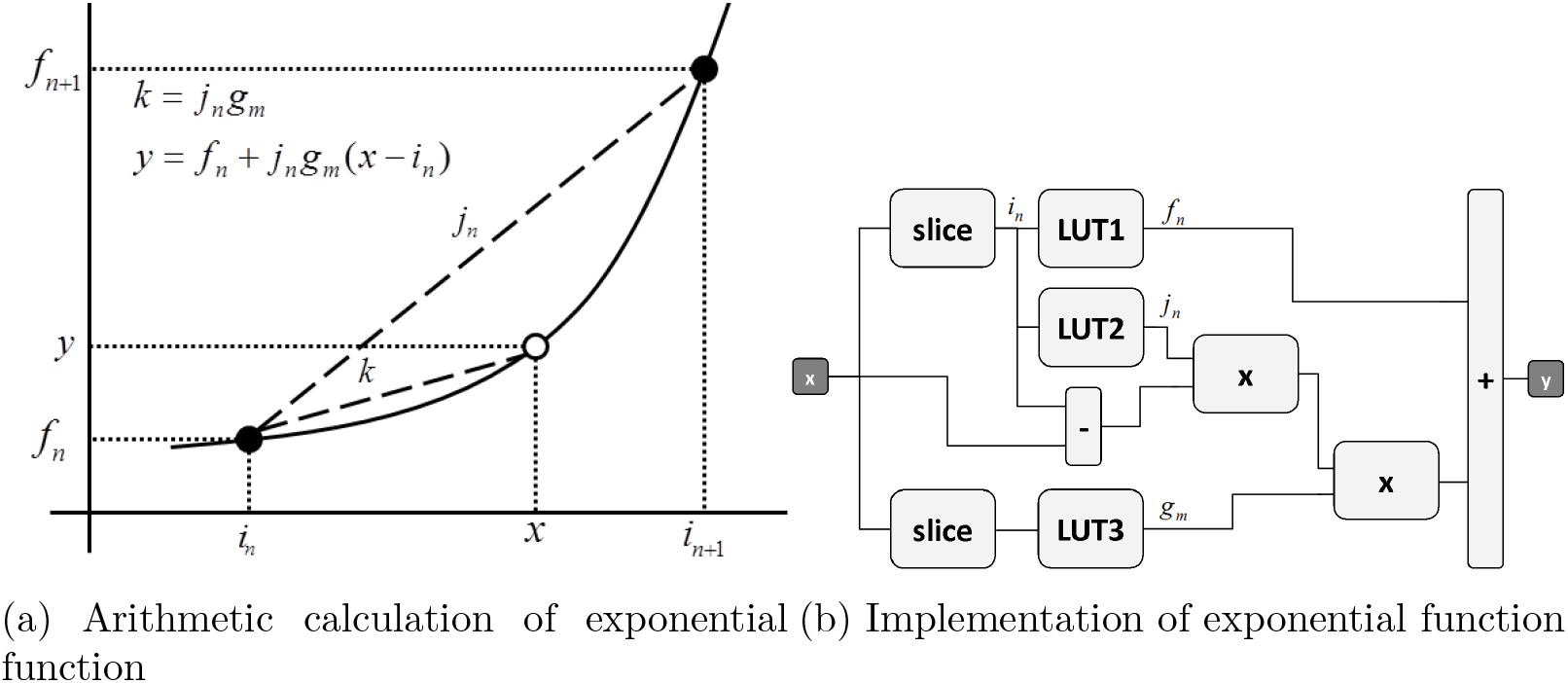
Design and architecture of the multipartite lookup-table exponential function implementation.

Through the use of dual-port memory devices the storage overhead can be reduced allowing for more efficient resource usage. Also, as the input values become more negative, the result of the exponential function tends to zero, further reducing storage requirements.

Table 4 shows a comparison between this technique, a pure LUT-based option and the optimized design presented by De Dinechin and Pasca (2010). As can be seen, the design presented here has reduced memory usage and an optimized latency.

**Table 4:** Comparison of Methods for Implementing Exponential.

|  | Our Design | LUT | De Dinechin and Pasca (2010) |
| --- | --- | --- | --- |
| Format | 28-bit Fixed Point |  | 32-bit Floating Point |
| Arithmetic | 2 Mult, 2 Add | 0 | 1 Mult, 3 Add |
| Memory | 16kbits | 15Mbits | 18kbits |
| Latency | 8 Clocks | 1 Clock | 15 Clocks |

### 3.3. Neuronal Network

For the development of large-scale neural models, that may incorporate thousands of neurons and synapses, a network-on-chip (NoC) approach to the FPGA neural design may be implemented. This could involve replicating the developed neural node multiple times across a single chip and attaching the node to a network router, whose responsibility it is to pass action potential information between different neurons within the network. This approach is illustrated within Fig. 6. This system could integrate the address-event representation (AER) scheme, whereby each neuron is allocated an address, and each spike produces a packet with this address attached to be broadcast (Furber and Brown, 2009). This is perhaps the most well defined protocol when considering neuromorphic network architectures and it has been used on many previous implementations (Boahen, 2000; Rast et al., 2011; Chicca et al., 2007).

**Figure 6:**
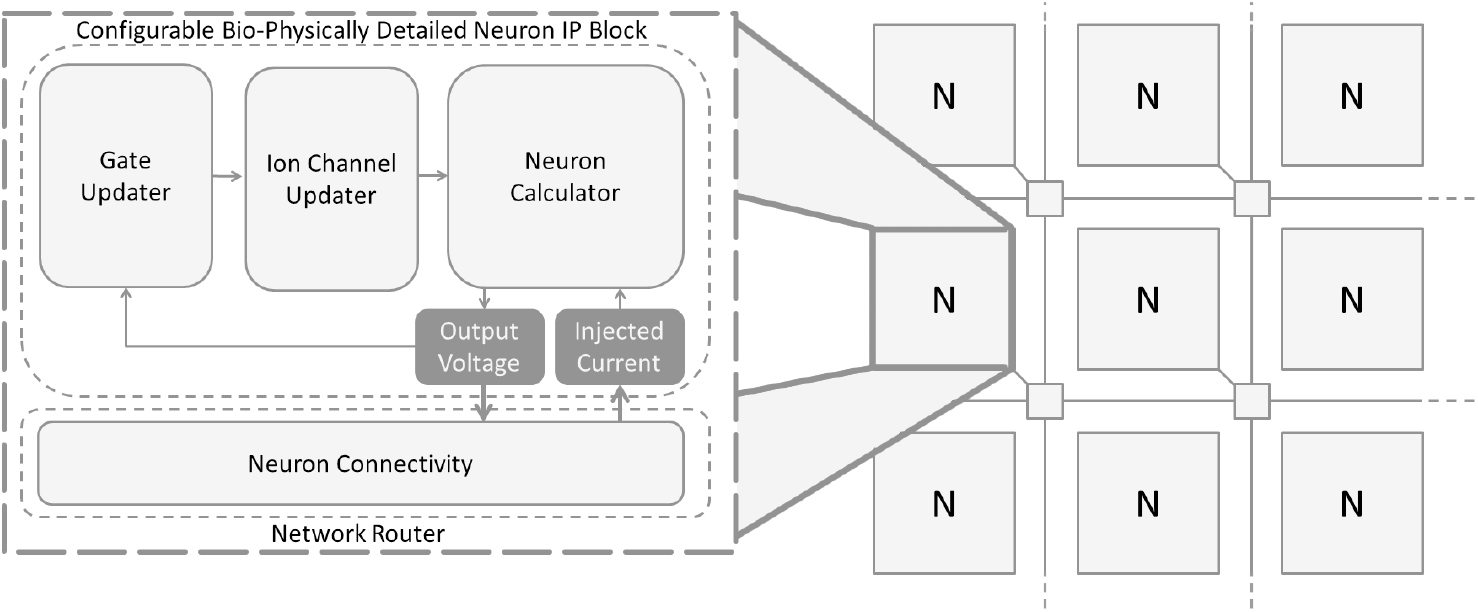
Large-scale network implementation. On a single FPGA multiple instances of the neuron IP block can be replicated. These instances then can be connected through a NoC infrastructure.

A better understanding of how damage to specific neural circuits affect brain function through reliable computer simulations boosts the speed and accuracy of diagnoses, and promotes the development of therapeutic treatments for neurological disorders or diseases. The mathematical models developed for the neural circuits also contribute to cutting-edge research in implantable neural prostheses, which aim to restore lost local nervous system function or enhancing brain function through closed-loop adaptive interactions with the brain in real-time. The hardware implementations of these mathematical models have already shown preliminary success in modulating simple intact neural circuits with mechanistic model or even more complicated regions in animals suffering neurological diseases or injuries using data-driven input-output model. The following section will describe such examples, which integrated computational emulation models and closed-loop feedback.

## 4. Hybrid Bio-Silicon Network

### 4.1. Dynamic Clamping Networks for Stomastogastric Ganglion

A dynamic clamp system (Sharp et al., 1993) consists of two main parts, the biological nervous system and the computational neuronal model. Fig. 7 shows an example of a basic configuration of a dynamic clamp system for interconnecting biological neuronal system with an FPGA.

**Figure 7:**
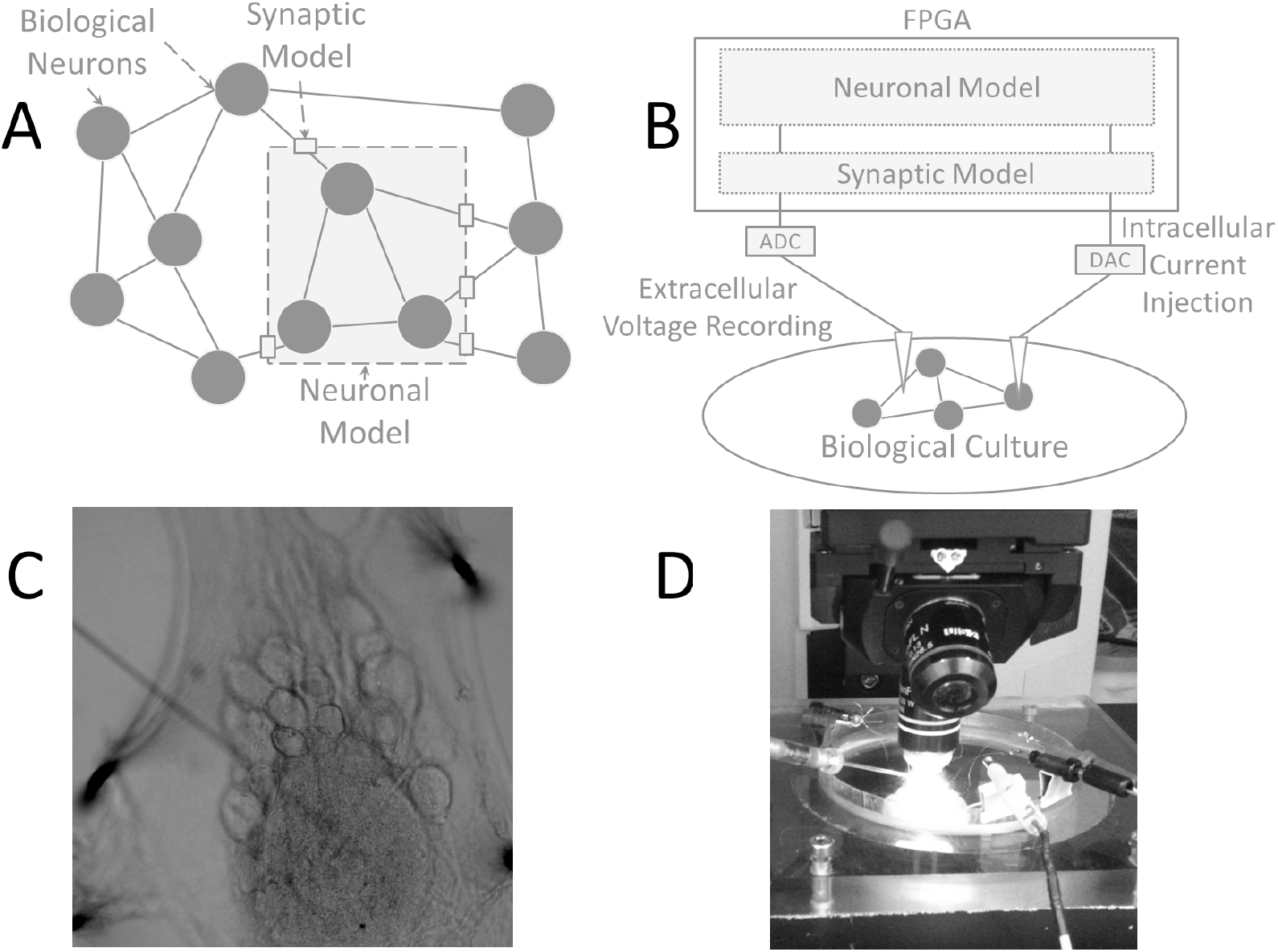
Hybrid network concept. A: Biological network may interact with a silicon neuronal model. B: The FPGA is able to interface with a biological culture through extracellular recordings and intracellular injection. C: An image of the biological culture, showing the STG nervous system and the intracellular glass electrode. D: Petri dish setup.

The implemented closed-loop system runs under recording mode and stimulation mode separately. The cell membrane voltage potentials are recorded by glass electrodes and are regarded as inputs into the computational model. The computational model is able to compute the feedback current and inject this to the recording cells.

For the biological experiments Adult Cancer Pagurus L. were obtained from local sources (Newcastle University, Dove Marine Laboratories) and kept in filtered seawater (10-12°C). Animals were kept in ice for 20-40 minutes for anesthetizing. We isolated the stomatogastric nervous system (STNS), and pinned this down in a silicone elastomer-lined (ELASTOSIL RT-601, Wacker, Munich, Germany) pet-i dish with chilled saline (10-13°C). The details of dissection and desheathing of the stomastogastric ganglion (STG) are followed from the article by Gutierrez and Grashow (2009). The neuron patterns generated in the STG are recorded using extracellular recordings.

The steps followed are as in our previous work (Luo et al., 2011). A petroleum jelly-based cylindrical compartment is built around a section of the main motor nerve (lvn) to electrically isolate the nerve from the bath; and one of two stainless steel electrode wires is placed in this compartment, the other one in the bath as a reference electrode. For intracellular recordings, current is injected using glass electrodes (GB 100TF-8P, Science Products, Hofheim, Germany; 25-40MOhm). A BA 01X Amplifier (NPI, Tamm, Germany) was used for signal amplifying. Files were converted and analyzed using Spike 2 v6.10 (Cambridge Electronic Design, Cambridge, UK).

In order to demonstrate the reliable function of the hybrid network setup, a system containing a biological cell connected to a model cell through a artificial synapse was created, this is the most common and useful application for dynamic clamp. Both electrical and chemical synapses were designed. For the electrical synapse, the architecture is followed as in Destexhe et al. (1994), where the activity of two neurons becomes synchronized. Whereas, for the artificial chemical synapse, the network displays half-center oscillations (Sharp et al., 1992)(Sharp et al., 1996).

Fig. 8 and Fig. 9 both show a significant positive correlation between the simulation and experimental results. Within Fig. 9 the pre-synaptic biological neuron reciprocally inhibits the post-synaptic neuron as expected. The silicon neuron burst frequency and inter-spike interval depend upon the inward current and chemical characteristics.

**Figure 8:**
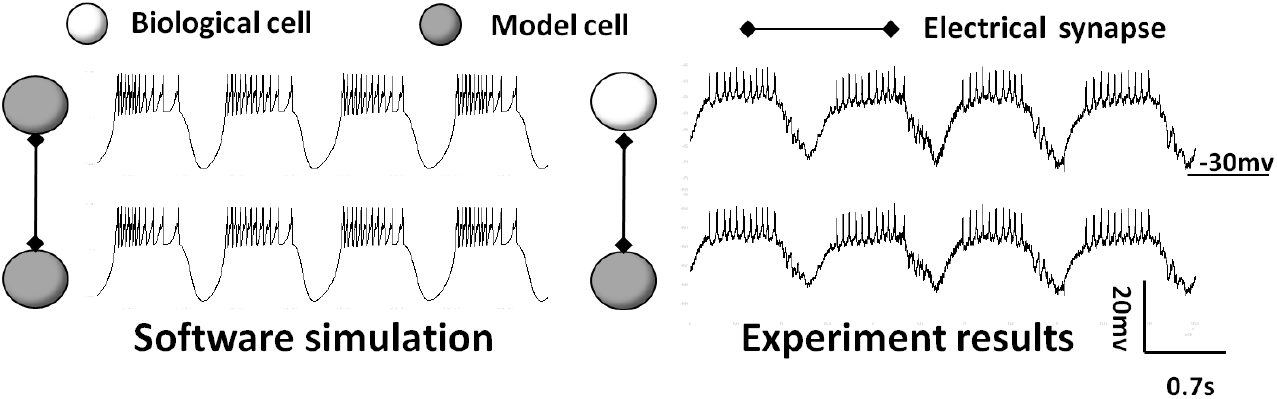
Electrical synapse simulation and biological results

**Figure 9:**
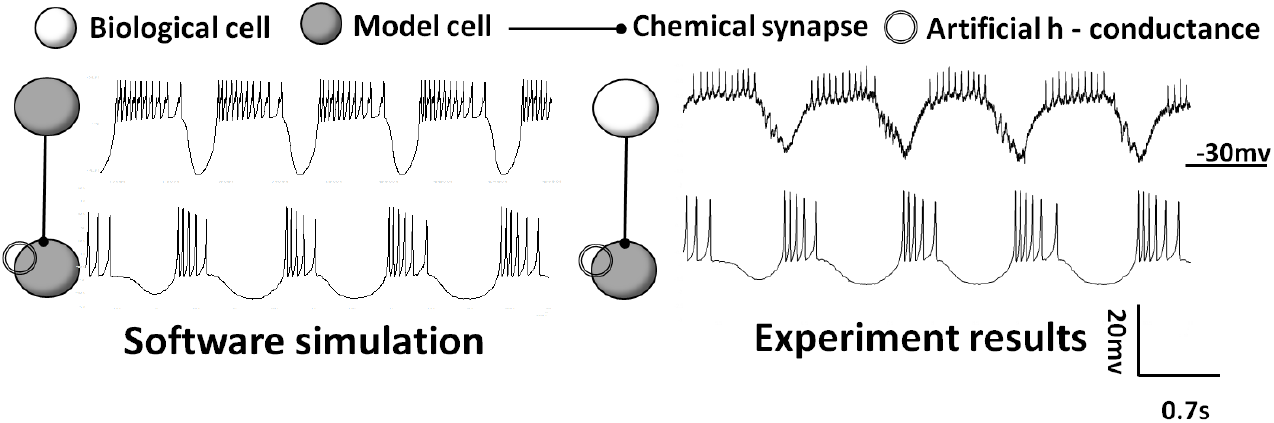
Chemical synapse interaction. The model cell reciprocally activates and deactivates with the biological cell

### 4.2. Hybrid Networks for Cellular Rehabilitation

The following hybrid network experiment employs STG neurons as the biological element. The experimental hypothesis is that the inactivated spatiotemporal dynamic of STG (AB and LP) neurons can be replaced by silicon neuron models, and that by reproducing the normal rhythm patterns and corresponding dynamics we confirm the system is reliable and practical. Results are shown in Fig. 10.

**Figure 10:**
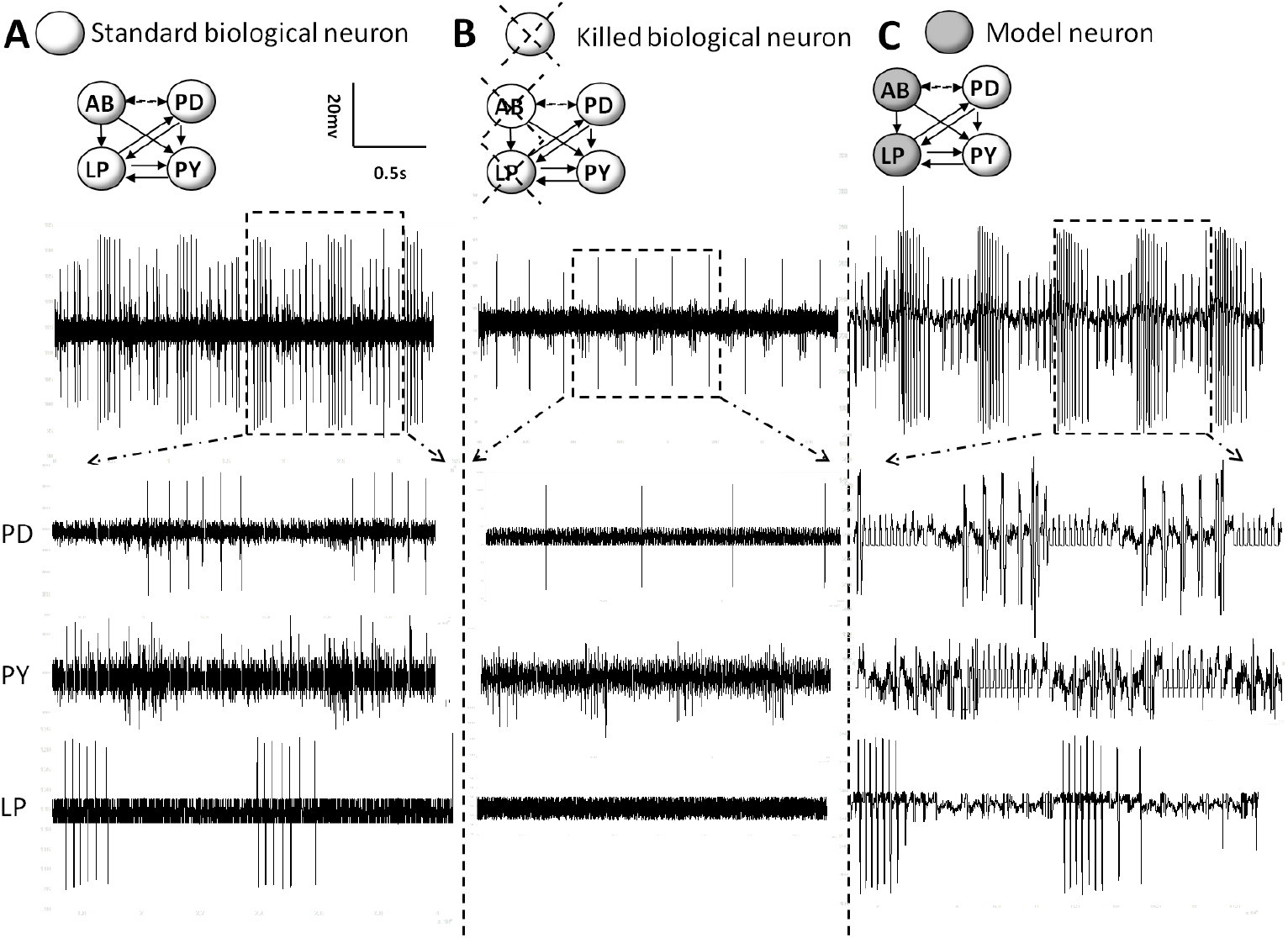
Results of hybrid bio-silicon network experiments. A: The intact biological network and its spike recordings. B: After two of the cells are deactivated the spike patterns are silenced. C: The patterns are rehabilitated by interconnecting the remaining neurons with silicon models.

Based on previous abundant research on the STG system, we have certain prior knowledge of the pyloric network: the AB neuron is responsible for generating rhythms in the network, and it reciprocally inhibits with the other neurons LP and PY. The other neurons are plateau neurons (Marder and Bucher, 2007; Selverston and Ayers, 2006; Nusbaum and Beenhakker, 2002). AB and LP neurons are deactivated to remove the pyloric rhythm.

The silicon neuron AB is designed as an intrinsic oscillator and LP has plateau neuron characteristics. The silicon synapses between AB and LP mimic the biological reciprocal synapse with artificial hyperpolarization conductance to display a half-centre oscillation network. The full interconnections between silicon and real cells are equivalent to the biological network topology. Model parameters are taken from Sharp et al. (1996), with slight modifications to fit the model architecture.

The results are shown in Fig. 10. In the normal network pattern (A), three different neurons generate individual patterns regularly and follow certain phase relationships. After deactivating neurons AB and LP, the network patterns have changed dramatically. PD shows single spikes irregularly, PY displays weaker patterns and LP is totally silent. One potential reason to explain this is that the AB neuron is the pacemaker of the pyloric network and therefore, it is responsible for generating network patterns and activating the other neurons.

Within section C of Fig. 10, two specifically designed silicon neuronal models AB and LP are interconnected with biological neuron, it is clear that the network rhythms are activated again; and they correspond relatively closely to the normal network rhythm.

We can quantitatively analyze this response using two measures, the burst period and duration for each recorded neuron. As shown by Fig. 11, the hybrid system is able to restore the burst duration of the PD neuron type, and the burst period and duration of the LP neuron. Minor modification of the system input parameters may reduce the slight variation between the hybrid and healthy scenarios in these measures. However, the results do demonstrate that the artificial neurons exert almost the same functionality in the biological network as the deactivated real cells AB and LP, indicating successful cellular rehabilitation of this neuronal network.

**Figure 11:**
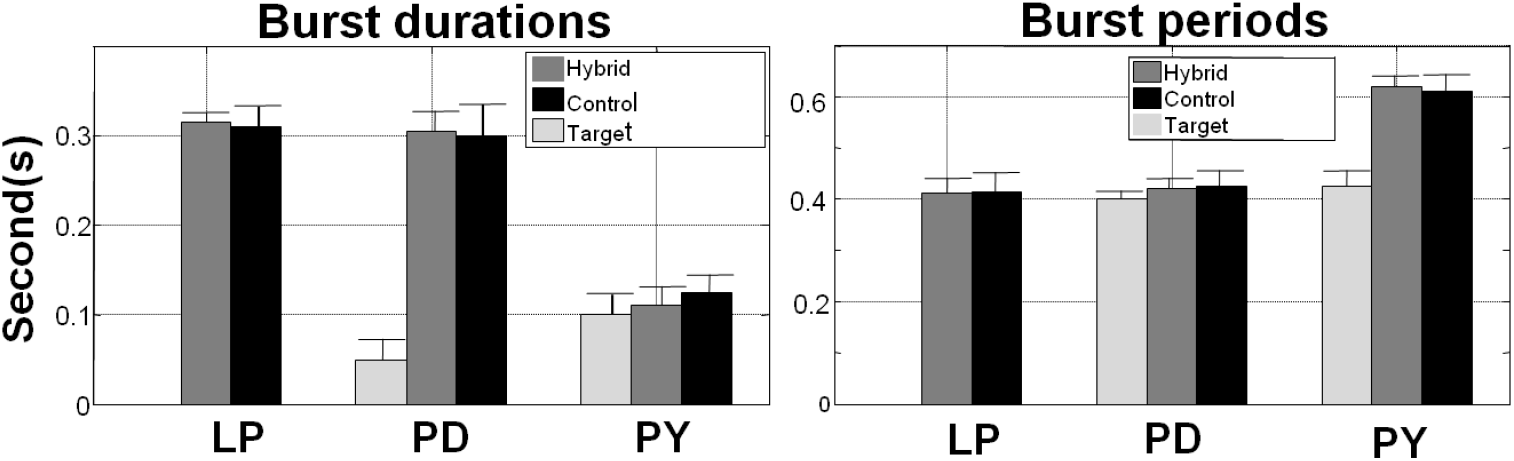
Comparison of burst periods and burst durations among unhealthy, healthy and hybrid networks

## 5. Discussion

### 5.1. FPGA-Hardware Utilization

The design of the ion channel simulator was created using the Xilinx System Generator software along with the Mathworks Matlab package. This is a common strategy used within other FPGA-based neural models (Luo et al., 2011; Weinstein et al., 2007; Mak et al., 2006; Graas et al., 2004). Initial simulations suggested a 28-bit signed fixed point system with a 14-bit fraction provided a sufficient level of resolution and range for most operations, although, the design requires minimal rework to alter the number system used. Within a dynamic clamping application the resolution of the input/output is constrained to between 14-16 bits depending upon the ADC/DAC used (Luo et al., 2011; Weinstein and Lee, 2006).

The design was synthesized for a Xilinx Virtex7 XC7V2000T FPGA using Xilinx XST. The key synthesis results are provided within Table 5. It can be seen that despite the addition of what was previously thought of as complex functionality, namely fixed-point exponential and division operations, the design uses only minor amounts of the available resources. After placing and routing the maximum clock frequency of the system was determined to be 240MHz. This allows for over 5000 ion channels to be updated within one integration time period using one ion channel block assuming an integration time step of 25*μs*.

**Table 5:** Resource usage comparison for different design approaches for ion channel simulation.

| Resource | This Work | Weinstein et al. (2007) | Mak et al. (2006) |
| --- | --- | --- | --- |
| Device | Virtex7 | Virtex4 | Virtex2 |
| Slices | 836 | - | 2800 |
| BRAM | 11 | 96 | 1 |
| DSP | 66 | 24 | - |
| Clock | 240MHz | 80MHz | 68MHz |
| Latency | 40 | 10 | 73 |
| Pipelined | Yes | Yes | No |
| Capacity | 256 Ion Channel Types | 6 Ion Channel Types | - |

Table 5 also provides estimated synthesis results for a LUT-based design. Although the latency and DSP usage is reduced in comparison to this design, as would be expected, the number of memory elements is substantially higher and the design only allows for 6 different compile-time derived ion channels to be utilized. The capacity in terms of ion channels of this work is only constrained by the size of the *Gate Types* ROM element, a doubling of this memory will result in double the amount of ion channel types.

Also shown within Table 5 are results from Mak et al. (2006) where the ion channel model used iterative calculations for the complex functions. The nature of this design breaks the pipeline and has a significant affect upon the latency of the calculation, thus reducing the overall throughput that would be achievable on a modern FPGA.

In order to simulate more ion channels it is possible to replicate the design multiple times across the FPGA, allowing for 150000 bio-realistic ion channels to be simulated on a single FPGA. This number is constrained by the DSP resource usage, currently at 3% of the overall resources available on a Virtex7 FPGA. This limits the number of times the design can be replicated. However, by reducing the DSP usage in a single ion channel block the number of channels that can be simulated will grow, as long as the fully pipelined status of the design and the clock frequency can be maintained. Alternatively, increasing the integration time period will also increase the number of channels, however this may cause instabilities in certain models. A comparison between the overall number of ion channels, and hence, the number of neurons that may be simulated on a single FPGA is provided by Table 6. As illustrated by this table, the design option described provides substantial rewards in terms of number of neurons that may be computed.

**Table 6:** Impact of resource usage on scalability of design assuming each design option may simulate 256 ion channel types.

| Design Option | Resource Usage/Node | Channels/Node | Channels/FPGA | Neurons/FPGA |
| --- | --- | --- | --- | --- |
| This Work | 3% | 5000 | 150000 | 50000 |
| Weinstein et al. (2007) | 100% | 5000 | 5000 | 1666 |
| Mak et al. (2006) | 1% | 83 | 8300 | 2766 |

### 5.2. Scalability

The importance of scalability when designing neuromorphic hardware is rapidly increasing due to the recent advancements in neural model complexity. As shown by Fig. 12a using a programmable approach offers significant rewards when the model complexity grows, due to the quadratic rises in memory requirements of the LUTs in an approach similar to Weinstein and Lee (2006). For each new type of gating variable introduced into the neural model the LUT-based approach requires a new block of memory to store the gating variable values. This increases the size of the channel simulator and therefore reduces the amount of times the simulator can be replicated across the whole FPGA, dramatically impacting upon the number of ion channels that can be simulated. However, a programmable approach is capable of calculating the values dynamically and hence there is no growth in resource usage as the number of gating variables or channel types grows.

**Figure 12:**
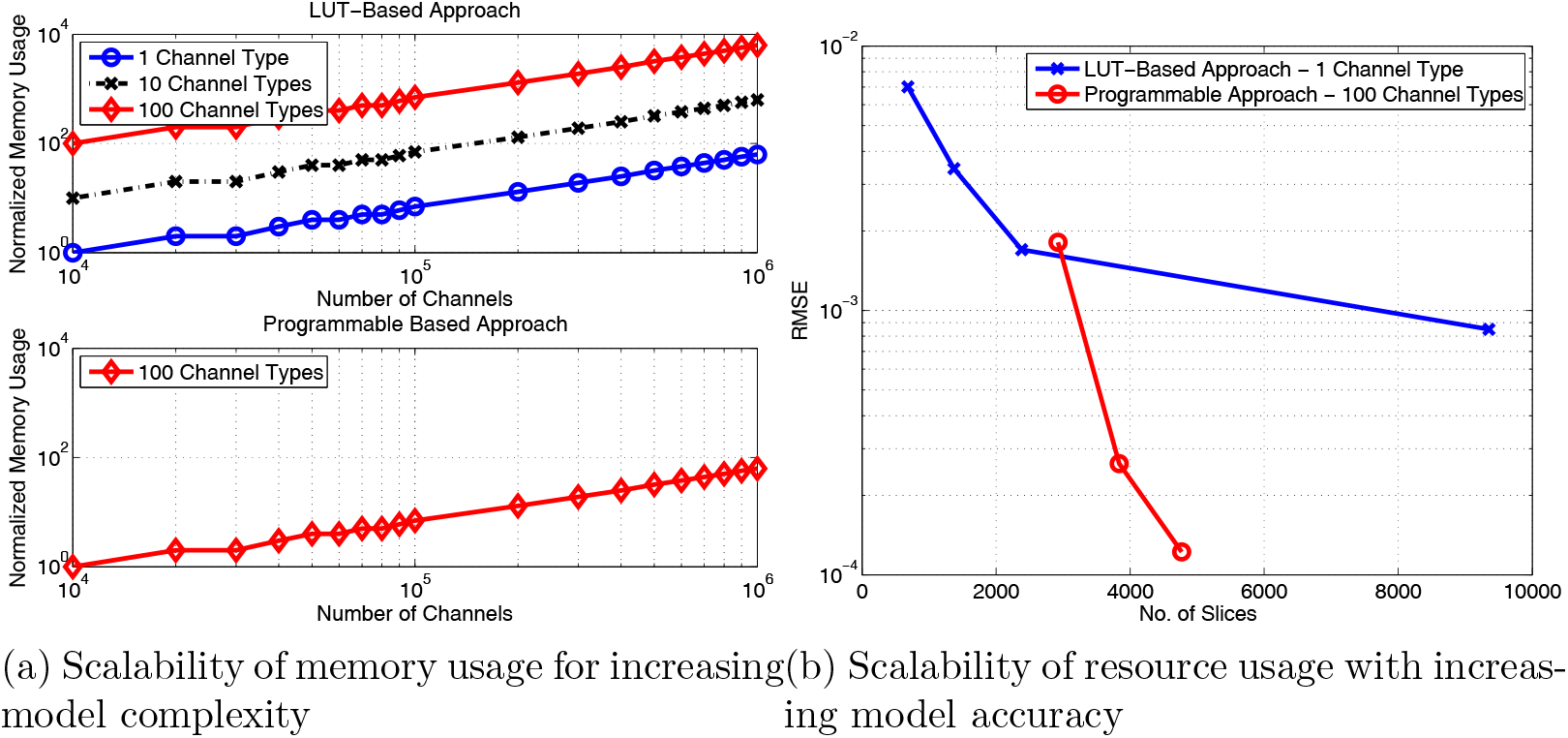
Design Scalability

Although the number system selected for this design gave adequate precision, it is easily configurable to add extra resolution if required. For example, due to the large number of ion channels that can be simulated the design could be utilized for simulations of detailed models of multi compartmental neurons (Graas et al., 2004), for which extra resolution may be beneficial. Fig. 12b shows how the resource usage of the two contrasting approaches to ion channel simulation scale with increasing resolution. In this figure all memory and DSP elements have been translated into slices. Using a pure LUT-based approach to reduce the error in the model by half the size of the LUTs must be doubled and this has a dramatic effect upon the overall number of slices. Whereas to reduce the error in a programmable approach the wordlength of the arithmetic calculations can be increased. Increasing the wordlength has less impact upon the number of slices used as opposed to doubling the size of the LUTs. Although the design offered by Mak et al. (2006) would scale in terms of area similarly to the approach offered here, the latency increased as the precision of the number system increased, due to the iterative nature of that design another iteration was required for each extra bit of resolution in the result.

### 5.3. Jitter, Reliability and Precision

Jitter is the variability in the actual duration of a computational cycle. For FPGA, jitter is barely existent in hardware-based dynamic clamp where a microprocessor is directly programmed. To test jitter issue in software and hardware, we compare variation of computation time of generating three bursts.

According to Fig. 13, it is very difficult to get stable and constant throughput using standard desktop operating systems. By contrast, FP-GAs still generate relative steady throughput while artificial neuron number increases. This reduction in jitter has important implications for developing large and complex network models.

**Figure 13:**
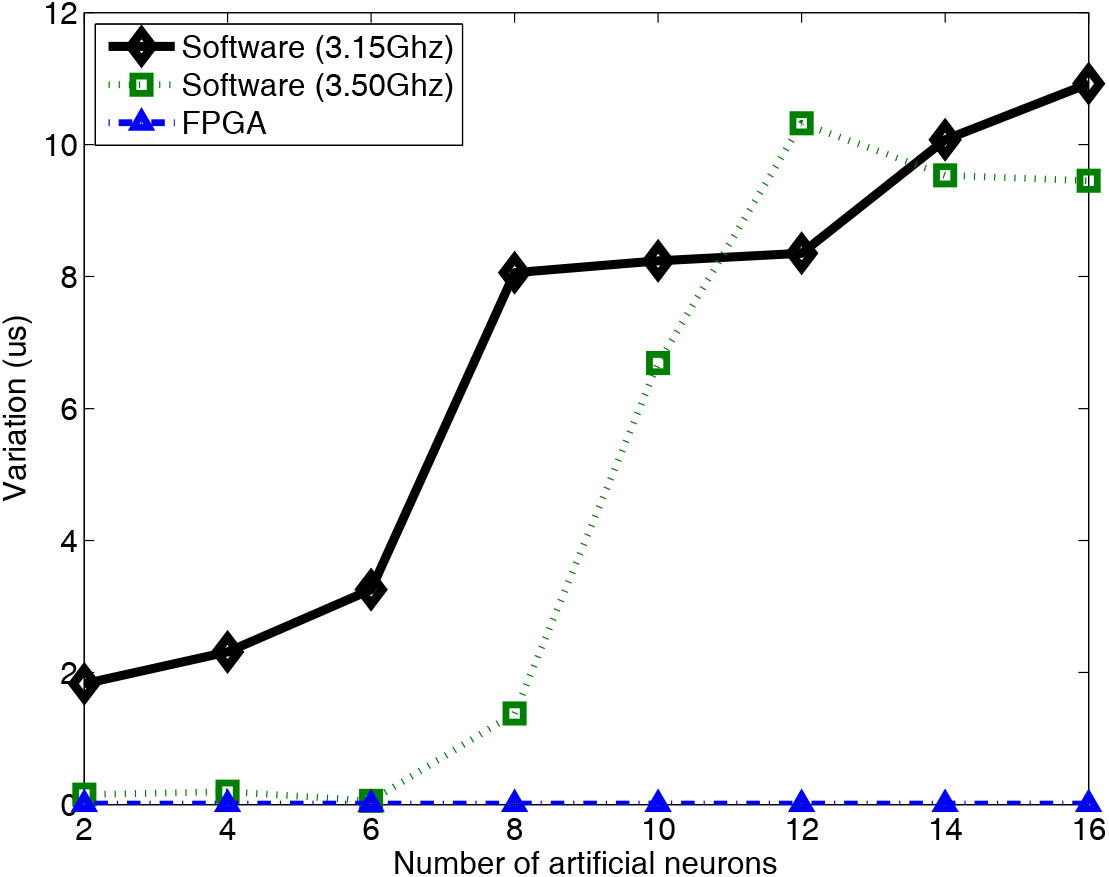
Comparison of jitter between software and FPGA

Fig. 14 shows the relationship between the spiking frequency and the injected current to a HH-type neuron. This is a common test of the stability of a neural system (Bettencourt et al., 2008) as the frequency and time stamp of the spike is thought to hold the key information. It can be seen that the dynamics of the low-precision FPGA-based design option corresponds closely with that of the high-precision software implementation.

**Figure 14:**
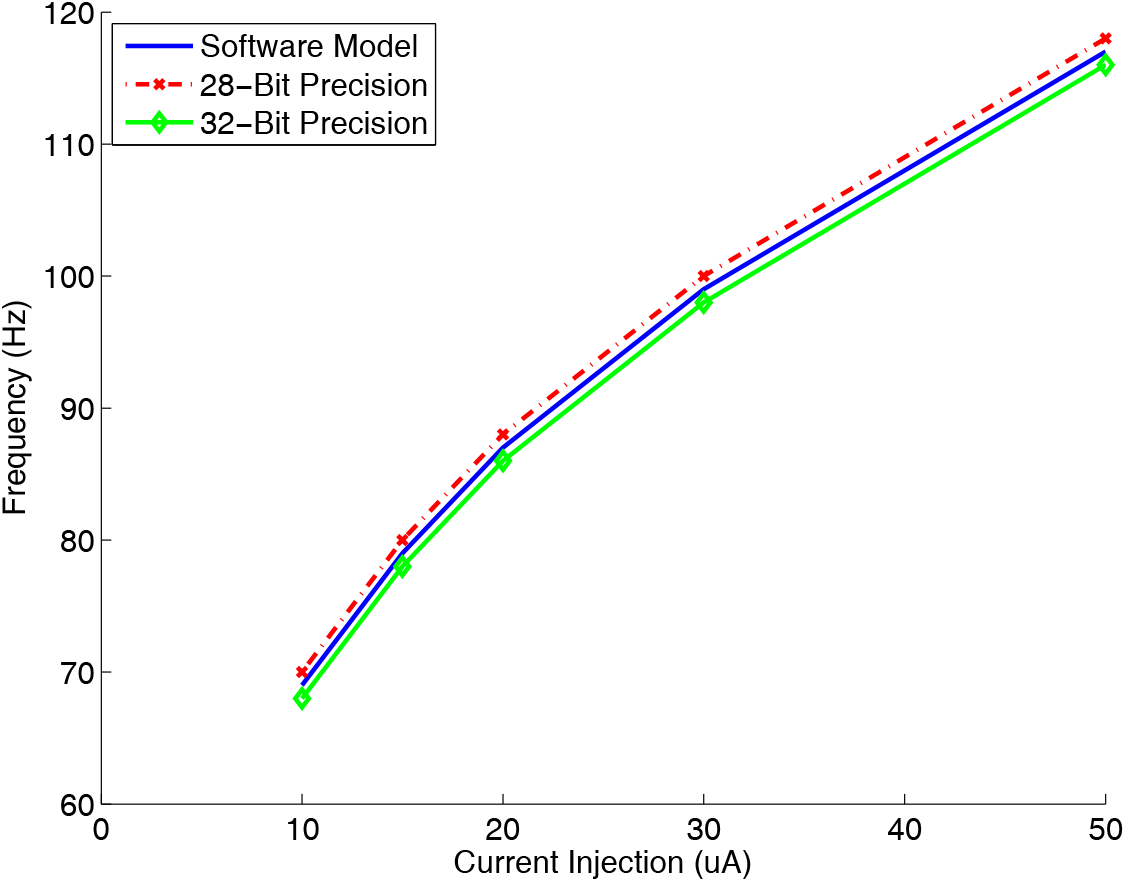
Spike frequency against current injection

## 6. Conclusion

The progress of BMI in recent years has been phenomenal, as demonstrated by the commercial success of cochlea prosthetics and the recent studies highlighting the potential for treatment for tetraplegia sufferers. Beyond these applications it is possible to foresee the development of cognitive prosthetic devices, which are capable of cellular rehabilitation, perhaps useful in conditions such as epilepsy.

Cellular prosthetic devices will rely heavily upon the formation of closed-loop hybrid networks, which consist of detailed silicon neuron models connected to biological elements through various neuro-silicon interfaces. The design of the silicon elements needs to be optimized for the task at hand, in order to provide the most efficient and powerful solution.

The implicit parallelism, reconfigurability and performance of FPGAs mean they offer a fantastic platform for the development of these nextgeneration silicon neural models, that are required to work in real-time and with high stability and reliability. As such, this paper has illustrated the design of an efficient and scalable complex neuron simulator. The simulations running upon this platform are able to be programmatically defined and it is capable of calculating over 50000 neurons on a single FPGA.

We have also illustrated the concept of cellular rehabilitation with practical biological experiments, involving a hybrid bio-FPGA setup. The functionality and rhythmic patterns of a small neuronal network were able to be recreated by reproducing the signals from deactivated neurons with silicon replicas. By implementing larger scale silicon neuron models the possibility of investigating the characteristics of more complex biological neural networks becomes a reality.

